# Insights into Gasdermin D activation from the crystal structure of its C-terminal domain

**DOI:** 10.1101/187211

**Authors:** Leonie Anton, Lorenzo Sborgi, Sebastian Hiller, Petr Broz, Timm Maier

**Author notes:** Corresponding author: Correspondence through Timm Maier at Department Biozentrum, University of Basel, Klingelbergstrasse 50/70, 4056 Basel, Switzerland.

## Abstract

Gasdermin D (GSDMD) is the central executioner of pyroptosis, a proinflammatory type of cell death. GSDMD is activated by the proinflammatory caspase-1 and caspase-11 via proteolytic cleavage in the linker connecting its N-terminal and C-terminal domain (GSDMD^Nterm^, ^Cterm^). The released N-terminal domain is sufficient to form pores in the plasma membrane, resulting in swelling and subsequent rupture of the cell. Here, we report the crystal structure of the autoinhibitory C-terminal domain of GSDMD at 2.04 Å resolution to further analyse determinants of GSDMD activation. GSDMD^Cterm^ adopts a compact helical fold unique to gasdermin proteins. Structural analysis and comparison to other gasdermin proteins reveals a conserved key interface for interactions between GSDMD^Nterm^ and GSDMD^Cterm^. Variations in two additional surface patches involved in interdomain interactions in full-length gasdermins suggest a role of these regions in modulating activation pathways, in agreement with biochemical characterization of different gasdermins.

## Introduction

Innate immune cells are part of the first line of host defense and recognize invading pathogens with the help of Pattern Recognition Receptors (PRRs) [1]. Membrane-bound PRRs monitor the extracellular or endosomal space for infection, while cytosolic receptors guard the intracellular space. PRRs recognize Pathogen-Associated Molecular Patterns (PAMPs) i.e. unique microbial molecules that are essential for survival or virulence. PAMPs are subject to low mutation rates and are typically conserved among different pathogens (e.g. Flagellin) [2]. In addition, PRRs also recognize Danger-Associated Molecular Patterns (DAMPs); endogenous host molecules, whose presence in a non-cognate compartment signals pathogen intrusion (e.g. ATP) [2]. PAMP/DAMP sensing by cytosolic receptors e.g. of the NOD-like (NLR) or PYHIN families [3, 4] triggers assembly of multi-protein signaling complexes, the inflammasomes (Fig. 1A). The first step in inflammasome assembly is PRR oligomerization, followed by recruitment of ASC (apoptosis associated speck-like protein containing a CARD) proteins. ASC recruits inactive pro-caspase-1 resulting in dimerization and autoproteolytic activation of caspase-1. Caspase-1 cleaves and matures proinflammatory cytokines (interleukin-1β and -18 (IL-1β/IL-18)), which are then released from the cell [5]. To release cytokines, caspase-1 activation also triggers pyroptosis, an immunologically active form of programmed cell death, mediated by a loss of integrity and ultimately lysis of the cell membrane. During pyroptosis, cells release DAMPs into the intracellular space, that are recognized by neighboring cells and trigger an inflammatory response [6, 7].

**Figure 1:**
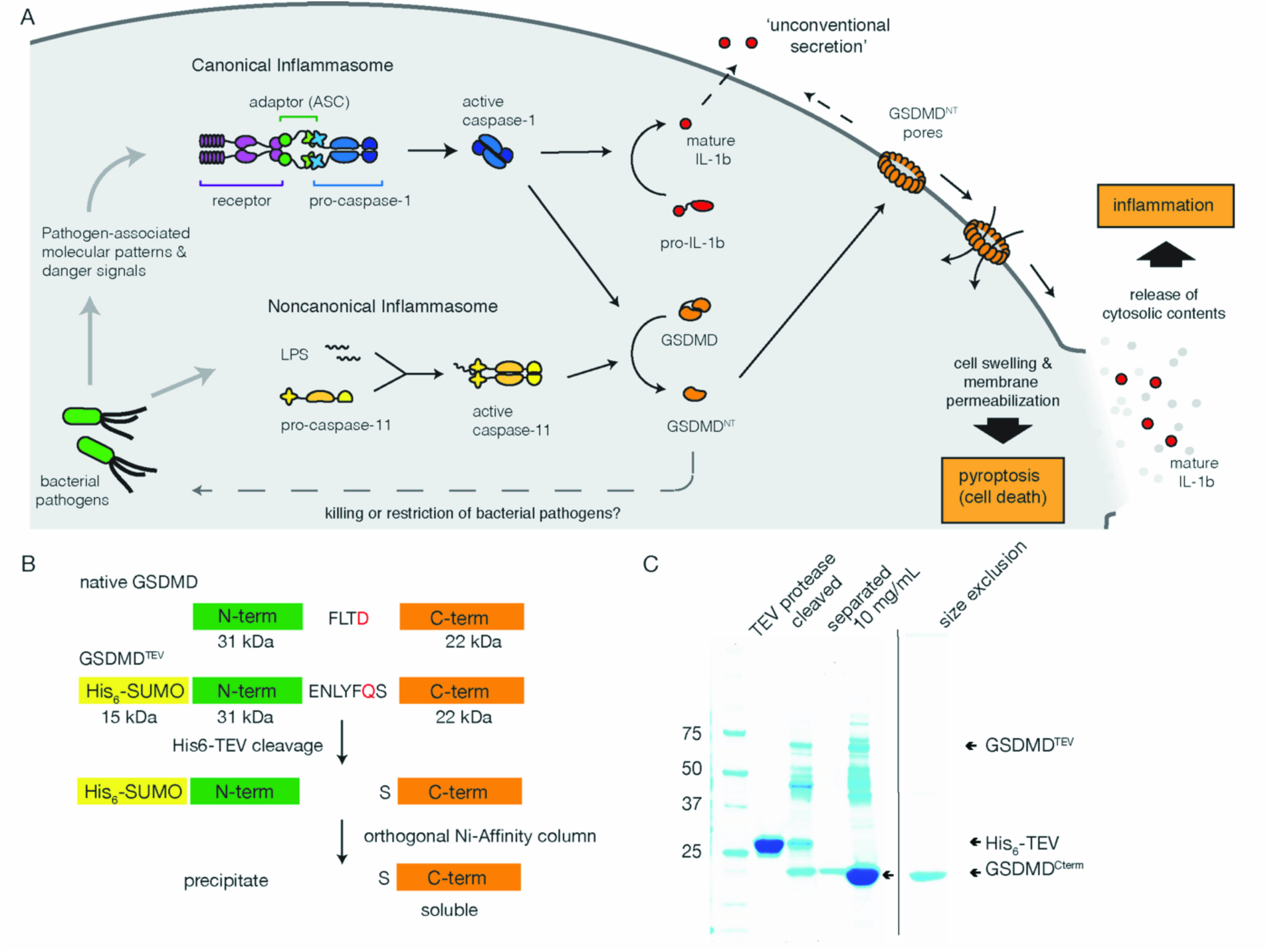
Efficient production of GSDMD^Cterm^ by *in vitro* proteolysis of GSDMD^TEV^. **A.** Schematic representation of *in vivo* GSDMD activation and function in inflammasome-dependent pyroptotic cell death. **B.** Domain architecture and protease cleavage sites of native GSDMD, GSDMD^TEV^ and overview of GSDMD^Cterm^ purification steps **C.** SDS-PAGE analysis of GSDMD^Cterm^ purification. Protein identity is indicated for GSDMD^Cterm^ (22 kDa), GSDMD^TEV^ (68 kDa), His_6_-TEV (28 kDa). TEV protease: TEV protease sample (1 mg/ml); cleaved: cleaved O/N sample; separated: sample after orthogonal Ni-affinity chromatography; 10 mg/ml: GSDMD^Cterm^ sample after concentration; size exclusion: peak fraction from size exclusion chromatography containing GSDMD^Cterm^.

Recently, the protein Gasdermin D (GSDMD) has been identified as the central executioner of pyroptosis. GSDMD is cleaved by caspase-1, but also caspase-11, the central protease in the non-canonical inflammasome [8, 9]. Human GSDMD is a 52 kDa protein composed of an N-and a C-terminal domain (hGSDMD^Nterm/C-term^) joined by a 44 amino acid (aa) linker. Active caspase-1 cleaves the hGSDMD linker at residue D275. Subsequently, GSDMD^Nterm^ forms irregular membrane pores and is necessary and sufficient for pyroptotic cell death [10-13].

GSDMD is a member of the gasdermin family consisting of six proteins in humans (hGSDMA, hGSDMB, hGSDMC, hGSDMD, hDFNA5, hDFNB59) and ten in mice (mGSDMA1-3, mGSDMC1-4, mGSDMD, mDFNA5, mDFNB59)[8]. Based on sequence similarity, all of them, except for DFNB59, adopt a similar two-domain architecture and are presumably activated by proteolytic cleavage. GSDMA, GSDMB, GSDMC, DFNA5 and mGSDMA3 induce pore formation in membranes upon artificial cleavage [10, 14]. In murine macrophages, DFNA5 is cleaved by caspase-3 and may promote secondary necrosis, a loss of membrane integrity in late apoptosis [14]. GSDMA and GSDMB polymorphisms have been implicated in asthma [15] and GSDMC mutations in tumorigenesis [16], but the native function and initiator proteases of these proteins are unknown.

The structure of the non-activated GSDMD^FL^ protein and the GSDMD^Nterm^ assembled into a functional pore are unknown. The available 1.9 Å crystal structure of murine full-length GSDMA3 (mGSDMA3^FL^) indicates a requirement for major conformational changes preceding pore formation in mGSDMA3^Nterm^, and – based on an overall sequence identity of 31.2% in the gasdermin family – presumably also in all related gasdermins including hGSDMD^FL^ [10]. Mutation of conserved aa in the hydrophobic interfaces between the GSDMA3^Nterm^ and GSDMA3^Cterm^ induce cell lysis similar to cleavage, suggesting an autoinhibitory function for C-terminal domain [10]. A recent crystal structure of hGSDMB^Cterm^ (25.1 % sequence identity with GSDMD) [17], rationalizes the effect of single nucleotide polymorphisms on hGSDMB function. The resulting mutations lead to decreased flexibility in a loop and increased positive charge, possibly interfering with the autoinhibitory function of hGSDMB^Cterm^.

To further define the mechanism of GSDMD activation in pyroptosis, we have determined a crystal structure of hGSDMD^Cterm^. Structural analysis and comparison to mGSDMA3^FL^ and hGSDMB^Cterm^ reveal together with additional biochemical data new perspectives on the activation of GSDMD and related gasdermins.

## Results and Discussion

### Design of an in vitro cleavable GSDMD construct

To obtain pure and soluble hGSDMD^Cterm^, the native hGSDMD caspase-1 cleavage motive _272_FLTD_275_, was replaced with a tobacco etch virus (TEV) protease cleavage site (ENLYFQ↓S, with cleavage occurring between Gln (Q) and Ser (S)) in the expression construct GSDMD^TEV^ (Fig. 1B). GSDMD^TEV^ was also N-terminally His_6_-SUMO-tagged, but the SUMO cleavage site was not used in the purification process (Fig. 1B). After purification of GSDMD^TEV^ by Ni-affinity chromatography, the protein was cleaved with His_6_-tagged TEV protease. His_6_-SUMO-GSDMD^Nterm^ precipitated upon cleavage. To separate the soluble GSDMD^Cterm^ from His_6_-TEV protease and uncleaved GSDMD^TEV^, an orthogonal Ni affinity chromatography step was performed. The GSDMD^Cterm^ protein product in the flow through was concentrated and further purified by size exclusion chromatography on superdex 75 (supp. Fig. 1A). Pooled fractions (Fig. 1C) were concentrated to 10 mg/ml for crystallization.

### GSDMD^Cterm^ adopts an all-alpha-helical fold characteristic of the gasdermin family

Purified GDSMD^Cterm^ was crystallized in the presence of sodium citrate and HEPES at pH 7 and 30°C yielding rod shaped crystals with dimension up to 20 x 120 μm. Coordinates of the C-terminal domain of mGSDMA3^FL^ were used to obtain phases by molecular replacement. In iterative cycles of manual model building and refinement, the model was refined to R-work/-free of 0.23/0.26 at a resolution of 2.04 Å in space group P2_1_2_1_2 (Table I).

**Table I.** Data collection and refinement statistics.

|  |  |
| --- | --- |
| Wavelength (Å) | 1.0 |
| Resolution range (Å) | 44.83 - 2.04 (2.113 - 2.04) |
| Space group | P 2<sub>1</sub> 2<sub>1</sub> 2 |
| Unit cell (Å) | 125.73, 47.98, 37.65 |
| $\alpha$, $\beta$, $\gamma$ (°) | 90, 90, 90 |
| Total reflections | 121073 |
| Unique reflections | 14421 (1432) |
| Multiplicity | 8.4 |
| Completeness (%) | 0.95 |
| Mean I/$\sigma$ (I) | 8.63 |
| Wilson B-factor (Å<sup>2</sup>) | 27.9 |
| R-meas | 19.6% |
| CC1/2 | 99.2 |
| Reflections used in refinement | 14419 (1432) |
| Reflections used for R-free | 749 (92) |
| R-work | 0.23 (0.30) |
| R-free | 0.26 (0.34) |
| Number of non-hydrogen atoms | 1656 |
| Macromolecules | 1495 |
| Protein residues | 199 |
| RMS (bonds) | 0.003 |
| RMS (angles) | 0.58 |
| Ramachandran favored (%) | 98 |
| Ramachandran allowed (%) | 1.5 |
| Ramachandran outliers (%) | 0 |
| Rotamer outliers (%) | 0 |
| Clashscore | 1.33 |
| Average B-factor (Å<sup>2</sup>) | 39.60 |
| Macromolecules | 38.91 |
| Solvent | 46.04 |
| Number of TLS groups | 7 |

Statistics for the highest-resolution shell are shown in parentheses.

GSDMD^Cterm^ adopts a compact all alpha-helical fold with ten alpha helices (α1- α10) and dimensions of 36.5 x 56.2 Å (Fig. 2A). The ten helices can be grouped into five N-terminal (α1- α5) and five C-terminal (α6- α10) helices, with α1, α2 and α5 interacting with α6, α7 and α10 to form the core of the protein. The N-terminal α1 and C-terminal α10 helices are aligned in antiparallel orientation and tightly interact via zipper-like hydrophobic interactions. The helices α2, α3 and α4 are connected by two long loops of fourteen and ten aa length, respectively, and form a helical bundle interacting with α5 and α6 (Fig. 2A).

**Figure 2:**
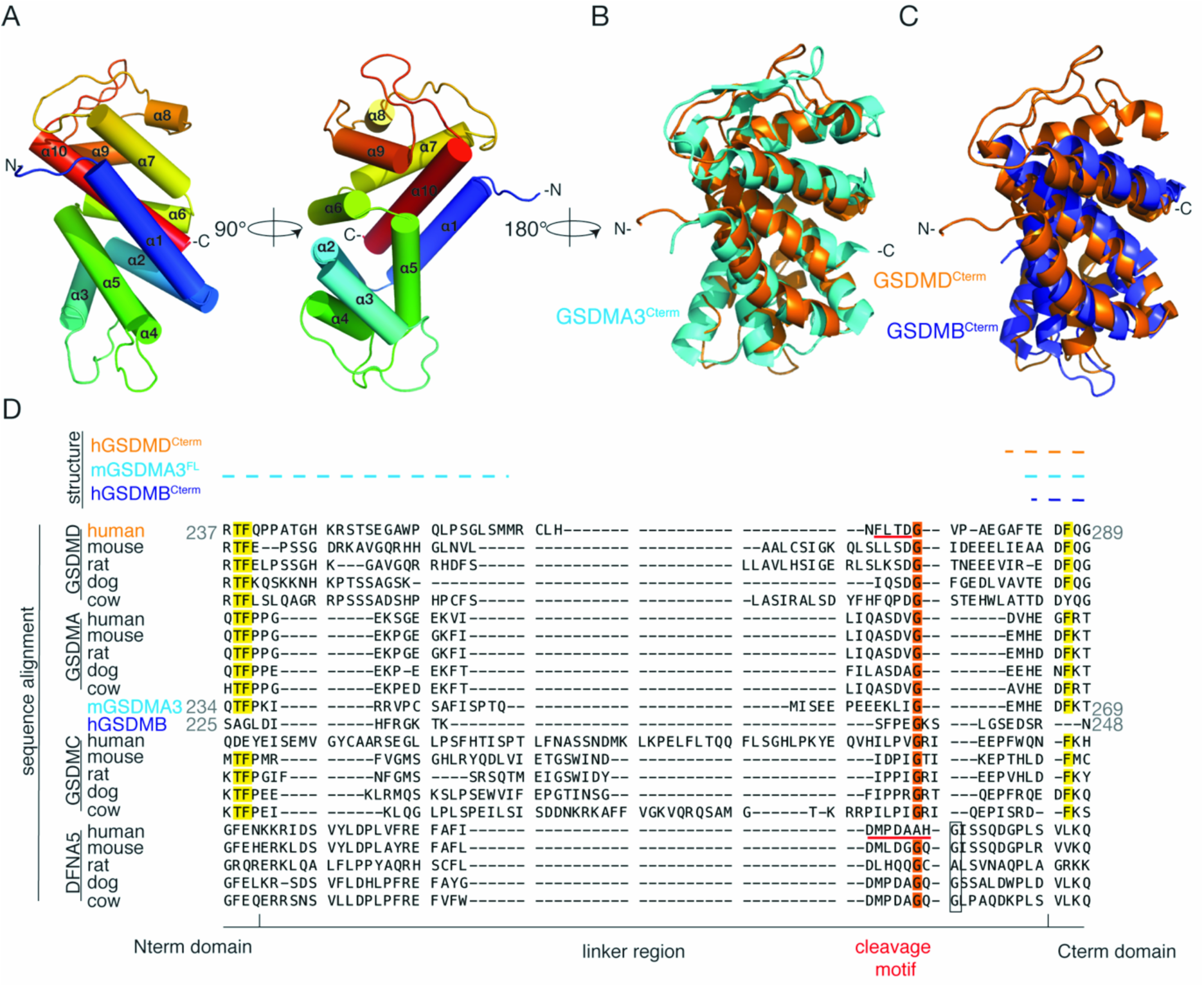
GSDMD^Cterm^ adopts an all-alpha helical fold characteristic of the gasdermin family. **A.** Cartoon representation of the crystal structure of hGSDMD^Cterm^. N-and C-termini are indicated and α-helices are shown as cylinders. superposition of hGSDMD^Cterm^ (orange) with mGSDMA3^Cterm^ (cyan, **B.**) or hGSDMB^Cterm^ (blue, **C.**). **D.** Sequence alignment of the linker region across different gasdermin families. Red: 100% conserved, orange: 80-100% conserved, yellow: 60-80% of conservation; bars at the top: sequence regions resolved in crystal structures of GSDMD^Cterm^, GSDMA3^FL^ and GSDMB^Cterm+MBP^ (color coded).

Secondary structure matching reveals that the fold of GSDMD^Cterm^ is unique to the C-terminal domain in the gasdermin family with a root mean square deviation of mainchain atom positions (RMSD) of 1.8 Å over 170 matched residues for mGSDMA3^FL^ (PDB:5B5R) at 32.9% sequence identity and an RMSD of 2.1Å for hGSDMB^Cterm^ fused to MBP (125 matched residues, 26.4 % sequence identity) (GSDMB^Cterm+MBP^, supp. Fig. 1B, PDB:5TIB) (supp. Table I).

Compared to mGSDMA3^Cterm^, hGSDMD^Cterm^ is characterized by an additional α-helix (α4), and shortened helices α3 and α9 (Fig. 2B, supp. Fig. 1C). hGSDMB^Cterm^ differs from hGSDMD^Cterm^ by a 34-residue deletion in the region of helices α8 and α9, an extension of helices α3 and α4 and a kink in helix α4 caused by a proline in position five of this helix (Fig. 2C, supp. Fig. 1D, supp. Fig. 2).

The best non-gasdermin family structural matches are with the PUB domains of human and murine peptide N-Glycanases (PNGase) (PDB:2CCQ/2HPL) with RMSDs of 3.1/3.4 Å over 80/86 residues matched to the α2, α4 helix and partially to α1, α5 of hGSDMD^Cterm^ at a sequence identity of 17.5/15.1% (supp. Table I, supp. Fig. 1E). The PUB domains of metazoan PNGases bind p97, which is involved in the unfolded protein response (UPR) and ubiquitin proteasome system [18], but their exact function is unknown. Gasdermins have so far not been implicated in related processes or interactions and the partial structural similarity is not sufficient to suggest a common function of gasdermins and PUB domains.

### The caspase cleavage site is embedded in a disordered interdomain linker

The caspase cleavage site in the interdomain linker of gasdermins is specific to their respective activation mechanism/enzyme and determines the first step in gasdermin activation [8-10]. Based on sequence and structural alignments, the linker extends from residue P242 to E285 in hGSDMD, equivalent to K234-E265 in mGSDMA3 and D225-R247 in hGSDMB (Fig. 2D). The length of the linker varies in gasdermins between 23 aa (hGSDMA) and 73 aa (hGSDMC) (Fig. 2D, supp. Fig. 2). The linkers in GSDMA and DFNA5 are predicted to have the same length (23/38 aa, respectively) across species, whereas linker length in GSDMD and GSDMC is variable (supp. Fig. 2). The highest variability of the linker region is observed when comparing GSDMDs across different species or to other gasdermins with only 22.5 % and 9.2 % sequence identity, respectively. Cleavage sites have so far only been experimentally identified for GSDMD (_272_FLTD_275_)(Fig. 2D, supp Fig. 3) and DFNA5 (_267_DMPD_AAH_273_) and are in equivalent regions of the linker [8, 14] (Fig. 2D). The Asp (D) of the caspase-1 cleavage site motif is strictly conserved in GSDMDs as well as the following Gly (G276 in human) (supp. Fig. 3). Cleavage sites are not conserved in other gasdermin families, however glycine residues in equivalent position of the linker are found in most gasdermins (Fig. 2D).

In the crystal structure of hGSDMD^Cterm^, the four C-terminal linker residues (A282, F283, T284, E285) have been modelled but are already characterized by increased flexibility. Weak electron density roughly representing four additional residues is visible, but could not be interpreted in terms of a unique peptide conformation. With four residues, this region would extend until P278 of the linker, just three residues upstream of the engineered TEV protease or two residues upstream of the native caspase-1 cleavage site (Fig. 2D). In the crystal structure of hGSDMB^Cterm+MBP^ only three aa (D245-R247) are modelled for the C-terminal part of the linker (PDB: 5TIB) [17]. The N-terminal region of the hGSDMB^Cterm+MBP^ linker is resolved in the structure, but is fused to a non-native MBP partner. It does not form internal interactions; its conformation is solely determined by contacts to non-native partners and is thus not of physiological relevance [17].

In the structure of the complete mGSDMA3^FL^, in the N-terminal part of the linker 15 aa after K234 and in the C-terminal linker region two aa (H263-D265) are ordered, while the central 15 residues remain disordered (PDB: 5B5R) [10] (Fig. 2D, supp. Fig. 2). The N-terminal part of the linker is folded into a loop held together by a short β-sheet between R236:T247 and R237:P246. This internal sheet structure of the linker may be of functional relevance, whereas the orientation of the loop is defined only by non-native crystal contacts [10]. The terminal residue of the N-terminal ordered linker region (Q248) is positioned by the internal sheet formation approximately 34 Å away from H264, the first ordered residue of the C-terminal linker part. This distance may well be spanned by the non-resolved 15 aa, however, only without the formation of larger helical secondary structures (supp. Fig. 1F).

In all three structures of hGSDMD^Cterm^, hGSDMB^Cterm+MBP^ and mGSDMA3^FL^, the ordered residues in the C-terminal part of the linker point in similar directions relative to the C-terminal domain (supp. Fig. 1G). In a structural superposition of hGSDMD^Cterm^ and mGSDMA3^FL^, one residue of the ordered part of the C-terminal linker region of hGSDMD^Cterm^ overlaps with the loop N16-D20 of mGSDMA3^Nterm^. This steric clash might be resolved in full-length GSDMD by a different structure of the non-conserved loop region in the N-terminal domain or by slight reorientation of the linker (supp. Fig. 1G).

Structural data on a full-length gasdermin family member including the linker region is currently only available for mGSDMA3, for which no proteolytic cleavage site has been identified, yet. However, the overall sequence similarity, the observed minimal length of the linker and the resolved structures of C-terminal linker fragments strongly suggest that all gasdermins contain a central highly flexible linker region comprising the proteolytic cleavage site.

### Partial conservation of interdomain interfaces in the gasdermin family

After caspase-1 cleavage, the GSDMD^Nterm^ and GSDMD^Cterm^ domains are no longer covalently linked and their interaction is defined only by protein-protein interfaces. The interactions between the N-and C-terminal domain have so far been only resolved for mGDSMA3^FL^ (PDB:5B5R) [10].

Sequence and fold conservation analysis shows a high degree of conservation in these interfaces, suggesting that the general interdomain architecture is conserved across the entire gasdermin family (Fig. 2B, Fig. 3A, supp. Fig. 2). Using published mutational data and sequence alignments we can thus define relevant interaction interfaces in hGSDMD^Cterm^ and their differences to other gasdermins. The calculated buried surface area (BSA) between the N-and C-terminal domain of mGSDMA3^FL^ is 2,684 Å^2^. In mGSDMA3, three interfaces between the N-and C-terminal domain are observed: The largest (Interface I, BSA 1,274 Å^2^) consists of a hydrophilic rim (light blue) with three hydrogen-bonds (hb) and two salt-bridges (sb) and a hydrophobic core (sky blue) (Fig. 3A, B), which is partially conserved in hGSDMD^Cterm^ (Fig. 3C). The two smaller ones, the hydrophilic interface II (1 hb, 2 sb, BSA 704 Å^2^), and interface III (no hb or sb, BSA 590 Å^2^) are less or not conserved, respectively (Fig. 3A, C, supp. Fig. 4).

**Figure 3:**
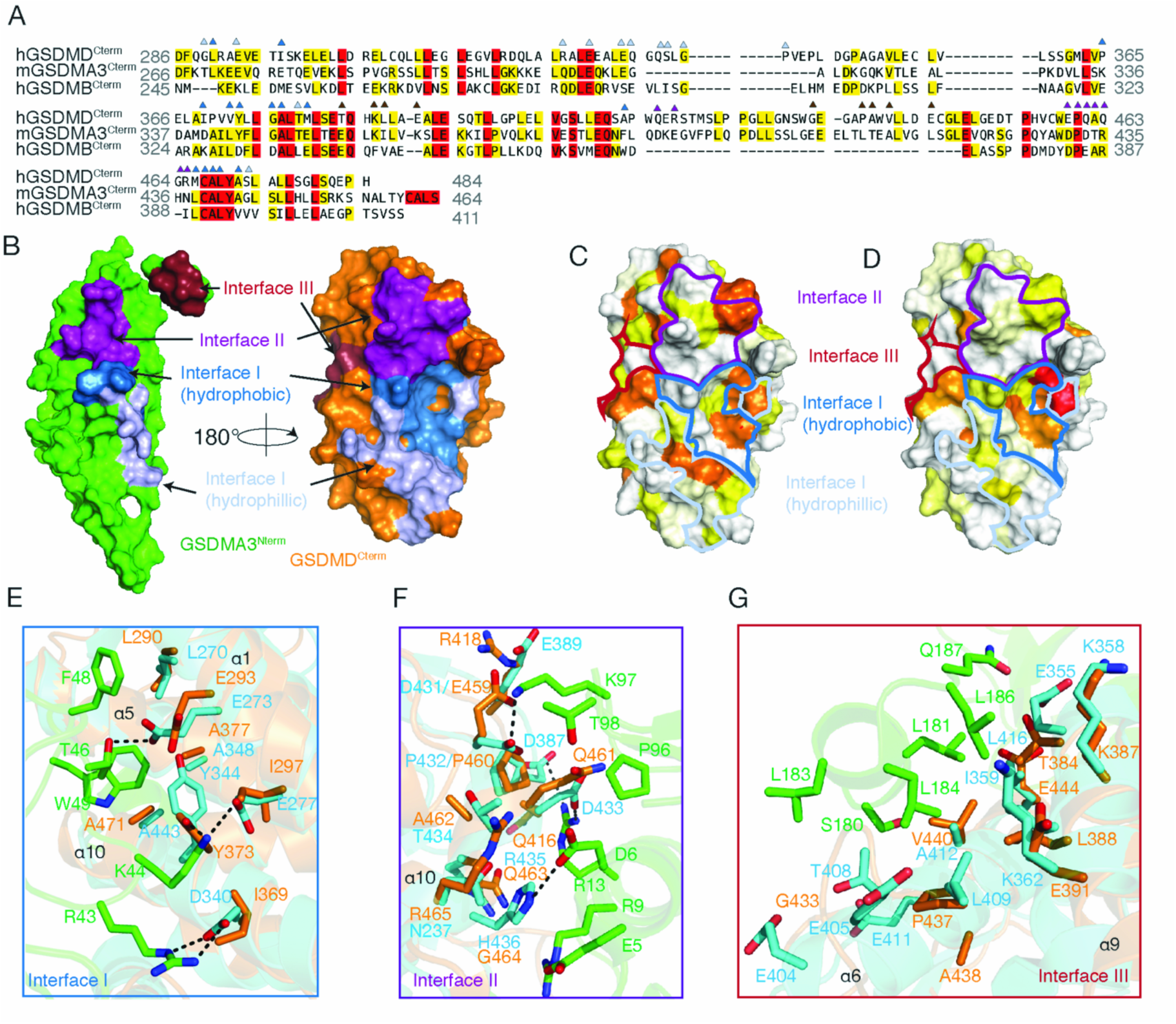
Analysis of interdomain interfaces in the gasdermin family. **A.** Sequence alignment of C-terminal domains of hGSDMD, mGSDMA3 and hGSDMB. Interface amino acids are indicated with triangles in colors corresponding to the interfaces (Interface I: hydrophobic: sky blue, hydrophilic: light blue; Interface II: purple; Interface III: dark red). **B.** Mapping of interfaces (color coded as in **A**) onto the surface of mGSDMA3^Nterm^ (green) and hGSDMD^Cterm^ (orange) structures. mGSDMA3^Nterm^ is rotated by 180° for better visibility. **C.** Mapping of conservation amongst GSDMDs C-terminal domains of different species and amongst all gasdermins onto the hGSDMD^Cterm^ structure (**D.**). Sequence identity amongst GSDMDs is 57.4 % and all gasdermins is 29.2 %. Color gradient indicates conservation between 60 % (red) and 0% (white). Interfaces are outlined in their respective colors. **E**- **G.** Close-up view of interdomain interactions in interfaces I (**E**, sky blue/light blue), II (**F**, purple), III (**G**, dark red) in GSDMA3^FL^.

The major interface I (Fig. 3B) in mGSDMA3^FL^ is formed by a loop of mGSDMA3^Nterm^, which inserts into a binding groove in the center of the interface in mGSDMA3^Cterm^. The loop (L40-V54) of mGSDMA3^Nterm^ includes six hydrophobic (L40, L47, F48, W49, G50, V54) and four polar or charged residues (K42, R43, K44, T46) involved in the interface. Two more polar residues (H146, K148) complete the hydrophilic rim around the hydrophobic core. These residues are conserved or conservatively substituted in hGSDMD^Nterm^, with the exception of only R43 (P44) and V54 (K55) (Fig. 3A, E). The binding groove for the loop in mGSDMA3^Cterm^ is formed by seven aa, which are located on the helices equivalent to α1, α3, α5 and α10 in hGSDMD^Cterm^ (Fig. 3E). Five of the seven aa in hGSDMD^Cterm^ (L290, E293, Y373, A377, A471) are strictly conserved between hGSDMD^Cterm^ and mGSDMA3^Cterm^. This part of interface I stands out as being highly conserved among GSDMDs and all gasdermins (Fig. 3A, C-E); and its particular relevance has previously been demonstrated: When transfecting HEK 293T cells with constructs expressing GSDMD with either of the residues L290, Y373 and A377 mutated to aspartate, cell survival decreases significantly compared to transfection with WT GSDMD [10]. Two of the groove forming aa are not conserved (E277, D340 in mGSDMA3; I297, I369 in hGSDMD) (Fig. 3E), but these are involved in the formation of three of the four hb in the center of the mGSMDA3^FL^ interface. The substitution will prevent formation of equivalent hb in hGSDMD, but might provide interface stabilization by hydrophobic interactions.

The rim of interface I is formed by nine polar residues surrounding the core groove in mGSDMA3^Cterm^. Four of these nine aa are conserved or conservatively substituted in hGSDMD (GSDMD/GSDMA3: G289/T269, R327/Q307, E330/E310, E334/E314, G340/D318, P341/K319, P365/K336, T379/T350, M380/E351) (Fig. 3A). The non-conserved interface II (purple in Fig. 3B) in mGSDMA3^FL^ is formed between a 3-residue loop (P96-T98) and the first helix of mGSDMA3^Nterm^ and a helix equivalent to a10 in hGSDMD^Cterm^ and comprises only hydrophilic interactions (Fig. 3F).

Interface III (dark red in Fig. 3B), is formed by hydrophobic interactions of short α-helix (S180-Q187) to the equivalent of helices α6 and α9 (Fig. 3G). Of the eleven aa involved in this interaction in the C-terminal domain, one is conserved (GSDMD/GSDMA3: K387/K358), five are conservatively exchanged (T384/E355, L398/I359, E391/K362, E434/E405, V440/A412), others are non-conserved (G433/E404, A436/T408, W439/E411, E444/L416, P437/L409).

Even though the overall structure of the C-terminal domain is very similar in hGSDMD^Cterm^, mGSDMA3^Cterm^ and hGSDMB^Cterm^, considerable variations in regions contributing to the predicted interfaces between the N-and C-terminal domains are observed. The structural comparison between hGSDMD^Cterm^ and mGSDMA3^Cterm^ demonstrates conservation of the central binding groove of interface I, while interfaces II and III are not conserved between GSDMD and GSDMA3. Interface I also shows the highest degree of conservation in GSDMDs of different species and amongst all gasdermins (Fig. 3C, D), as supported by published mutational data [10]. The remaining interfaces vary in their degree of conservation and are presumably responsible for modulating interdomain interactions in different gasdermins.

### Post-Cleavage Activation in different GSDMs

The presence or absence of inherent interactions between the N-and C-terminal domain of gasdermins determine the post-cleavage activation steps. The structural differences observed in the interdomain interfaces, open up the possibility for gasdermins to have distinct post-cleavage behavior tuned for their respective functions.

Cleavage of hGSDMD^FL^ leads to the direct dissociation of the N-and C-terminal domain. In the presence of membranes, hGSDMD^Nterm^ spontaneously forms membrane pores as we have recently demonstrated [12], while in the absence of lipids it precipitates. This precipitation of hGSDMD^Nterm^ might be the results of conformational changes or the missing stabilization by hGSDMD^Cterm^. Although these results indicate a hGSDMD^Cterm^ independent downstream activation of hGSDMD^Nterm^, overexpression of isolated hGSDMD^Cterm^ in HeLa cells inhibits pyroptotic cell death [8] induced by hGSDMD activation.

Contrastingly, Ding et al. recently reported for mGSDMA3, that after linker cleavage non-covalent interactions prevent dissociation of the N-and C-terminal domains [10]. Final execution of physiological function would thus likely require an additional activating step for dissociation of the N-and C-terminal domain.

For human GSDMA, we and others note that after cleavage by TEV protease, the two domains dissociate as demonstrated here by chromatographic separation (Fig. 4A, supp. Fig. 1B) [10]. Remarkably, no precipitation of hGSDMA^Nterm^ is observed after cleavage (Fig. 4B) indicating that in contrast to hGSDMD, hGSDMA^Nterm^ retains a certain solubility even in the absence of GSDMA^Cterm^. In the presence of plasma membrane-like liposomes, hGSDMA^Nterm^ forms pores, but this process is less efficient than for hGSDMD and mGSDMA3 [10]. Only in cardiolipin-containing liposomes, hGSDMA forms pores with similar kinetics as compared to hGSDMD and mGSDMA3, providing evidence for an effect of lipid composition of membranes on pore formation by N-terminal domains of gasdermin proteins (Fig. 4C).

**Figure 4:**
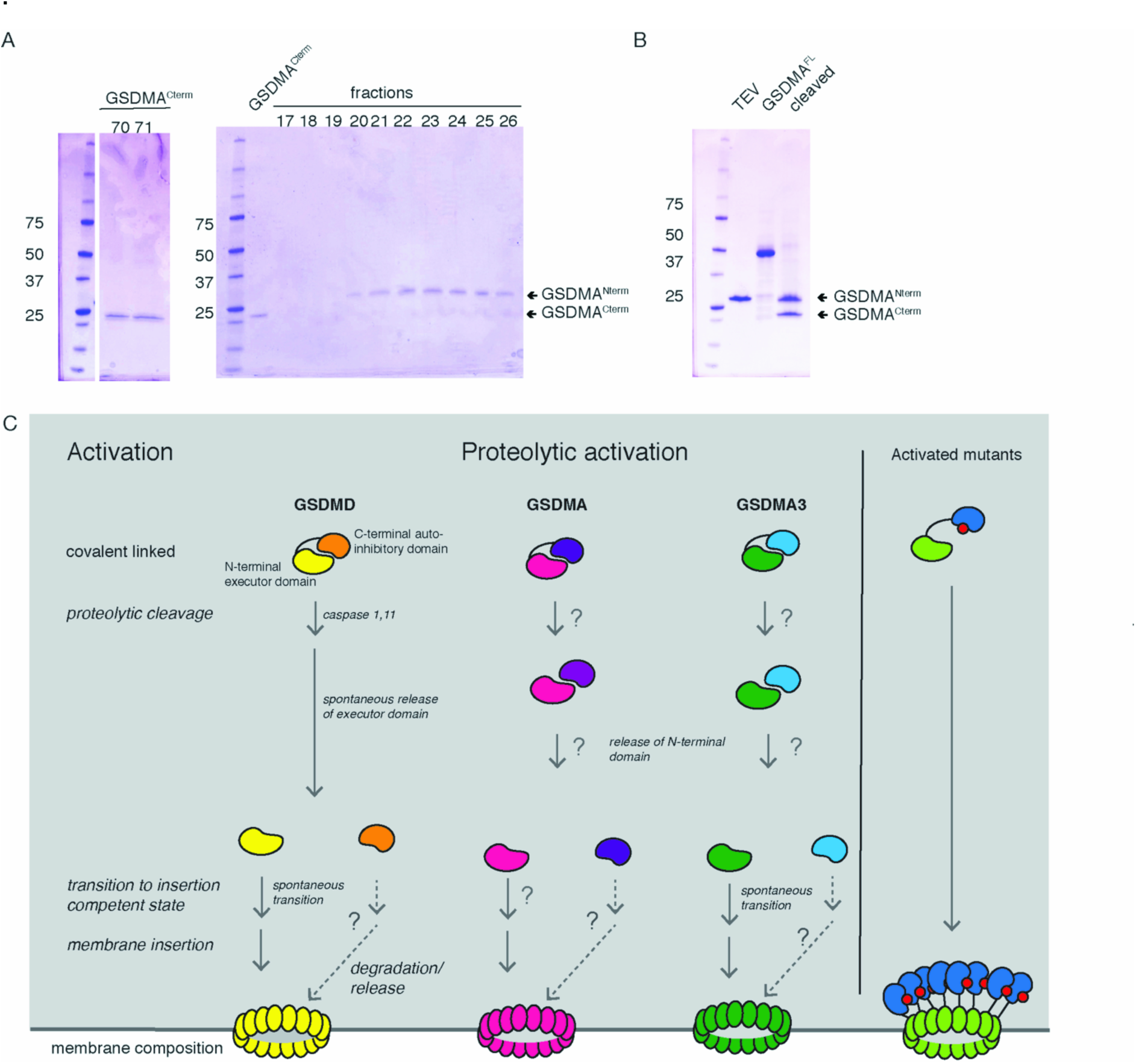
Activation of gasdermins is a multi-step process. **A.** GSDMA^Nterm^ (27 kDa) and GSDMA^Cterm^ (23 kDa) are separated by anion exchange chromatography. TEV: TEV-protease (1 mg/ml) **B.** After 3h cleavage of GSDMA^FL^ (50 kDa) with TEV (28 kDa) two bands are visible (21 kDa, 27 kDa). **C.** Schematic representation of gasdermin activation via proteolytic cleavage, transition to a membrane-insertion competent state and finally membrane insertion and pore formation (left panel, for details see text). Membrane composition may influence pore formation. The right panel illustrates that mutations in the interdomain interface may be sufficient to release autoinhibition by the C-terminal domain.

The dissociation of the N-and C-terminal gasdermin domains depends on their specific non-covalent interactions and possibly additional interacting factors (Fig. 4C). Based on the observed precipitation, hGSDMD^Nterm^ is not efficiently stabilized by hGSDMD^Cterm^, possibly due to a weaker interaction of the two domains or a higher potential of hGSDMD^Nterm^ for conformational transitions towards an insertion competent state (Fig. 4C). Altogether, these observations indicate that gasdermins, despite sharing pronounced structure and sequence similarity, display different post-cleavage behavior *in vitro*

Here, we determined the crystal structure of hGSDMD^Cterm^, responsible for auto-inhibition of the executioner of pyroptosis, hGSDMD [8, 9], and analyzed the partly conserved role of the C-terminal domain in gasdermins. The first step in the activation of GSDMD is the proteolytic cleavage by caspase-1 within the central linker region, which is disordered around the caspase recognition site and not conserved among gasdermins. The C-terminal domains of gasdermins share high sequence similarity and adopt a unique conserved helical fold. Based on the structure of mGSDMA3^FL^, we analyze plausible interactions between N-and C-terminal domains of GSDMD and other gasdermins and identify at the structural level – in agreement with previous mutational studies [10] — a conserved core interdomain interaction site. Additional non-conserved interfaces modulate the interactions between the N-and C-terminal domains resulting in differential dissociation, as showcased by the difference in post-cleavage behavior of hGSDMD, mGSDMA3 and hGSDMA.

Further research will be required to identify the proteases responsible for the cleavage of the different gasdermin family members and their respective activation pathways. Our current work indicates that the final execution of function might as well require specific environments or activating interactions beyond proteolytic cleavage, both for domain separation and for promoting membrane insertion of the N-terminal executor domains in some gasdermin family members.

## Materials and Methods

### Protein Expression and Purification

cDNA coding for the full length human GSDMD was cloned with an N-terminal His6-SUMO-tag into a pET28a vector under control of a T7 promoter. TEV protease cleavage site was incorporated by QuickChange site-directed mutagenesis kit (Stratagene, Agilent Technologies, La Jolla, California) to create GSDMD^TEV^. All plasmids were verified by DNA sequencing. Protein expression and purification were done as described in Sborgi et al [12]. Protein constructs were transformed in BL21 (DE3) *E. coli* strains, and the proteins were expressed by growing the cultures at 37°C to an OD600 of 0.7 and by inducing with 0.5 mM IPTG overnight at 18°C. The cells were harvested by centrifugation and the pellet was resuspended in 20 mM Tris, pH 7.5, 50 mM NaCl, 5 mM imidazole, 20 mM MgCl2, 10 mM KCl, 0.5 mM TCEP, 0.1 mM protease inhibitor mix (PMSF, Pepstatin, Bestatin, Phenanthrolin, Leupeptin), and DNase I. Resuspended cells were disrupted by microfluidization and centrifuged at 30,000 g at 4°C for 45 min. The supernatant was filtered and loaded onto a Ni-NTA affinity column (Genscript, George Town, Cayman Islands). The column was washed with 20 column volumes (CV) of resuspension buffer containing 15 mM imidazole, and the fusion protein was eluted with 5 CV of the same buffer with 250 mM imidazole. GSDMD^TEV^ was cleaved by His_6_-tagged TEV protease O/N at 4°C or for 4 h at RT, respectively. GSDMD^Cterm^ was collected from the flow through from an orthogonal Ni-NTA affinity column step and concentrated. Using a superdex 75 gel filtration column (GE Healthcare, Chicago, Illinois) pre-equilibrated with 20 mM Tris buffer pH 7.5, 50 mM NaCl, 0.5 mM TCEP, monomeric GSDMD^Cterm^ was collected in fractions and pooled. The sample was concentrated to 10 mg/ml and was frozen in small aliquots in liquid nitrogen. A similar purification protocol was applied to GSDMA^TEV^ with the following modifications. A TEV cleavage site was generated using a PCR with overlapping primers. GSDMA^TEV^ was cleaved with ULP1 protease O/N at 4° C to cleave off the His_6_-tag. The GSDMA^Cterm^ and GSDMA^Nterm^, which are in a non-covalent complex after cleavage by His_6_-TEV protease, were subjected to another orthogonal Ni-NTA affinity step and concentrated. The two domains were separated using a 10x100mm PEEK anion exchange column with POROS HQ/M20 resin (Thermo Fisher Scientific, Waltham, Massachusetts) equilibrated in 0.02 M Tris pH 7.5, 0.05 M NaCl and eluted with a gradient of 0.02 M Tris pH 7.5, 1 M NaCl. All samples were analyzed using SDS-PAGE.

### Crystallization and Crystallographic Data Collection

All crystallization experiments were conducted using sitting drop vapour diffusion. Crystals were grown at 30°C by mixing protein and reservoir solution (1.229 M Sodium Citrate, HEPES, pH 7.25) in a 1:1 or 2:1 ratio. Thin rod shaped crystals appeared on day one and continued growing until day five. Crystals were cryoprotected using cryo-oil (Perfluoropolyther Cryo Oil, Hampton Research, Aliso Viejo, California) and flash-cooled in liquid nitrogen. Crystallographic data were collected at the Swiss Light Source (Paul Scherrer Institute, Villigen, Switzerland) at beamline X06SA at 100 K using a Eiger 16M detector (Dectris, Baden-Daettwil, Switzerland). GSDMD^Cterm^ data were collected at a wavelength of 0.999998 Å with an exposure time of 0.02 s, a rotation angle of 0.25°, and a detector distance of 0.225 m.

### Structure Determination of GSDMD^Cterm^

GSDMD^Cterm^ data were processed using XDS [19, 20]. Phases were obtained by molecular replacement using PHASER [21]. The initial model was refined by iterative cycles of manual model building in Coot [22], density improvement in parrot [23], model autobuilding in buccaneer [24, 25], and refinement in buster [26] and phenix [27-30]. The final model includes residues 282 to 480 with residues 282-285, 299-307, 334-344 and 360-367 showing increased disorder of side-chains. 278-281 could not be modeled due to pronounced disorder. Structure factors and model were deposited and approved by PDB under ID 5NH1.

### Structural Analysis

Secondary structure matching was done using PDBeFold, with parameters for query structure at 80 % lowest acceptable match and 40 % lowest acceptable match for target structure [31] or direct comparison to PDB structures of mGSDMA3^FL^ (PDB:5B5R) and hGSDMB^Cterm+MBP^ (PDB: 5TIB). Interfaces of proteins were analyzed by PISA [32]; alignments of sequences were done using Clustal Omega [33, 34] and manually optimized. Conservation was plotted using AL2CO, with the entropy-based algorithm [35] and figures were generated using the PyMOL Molecular Graphics System, Version 1.7 (Schrödinger).

## Acknowledgements

We would like to thank the staff of the PX1 beamline at the Swiss Light Source in Villigen (CH) for their excellent support.

## supplemental material

Supplemental materials include additional materials on sequence and structure-based comparison of different members of the gasdermin protein family, structural superpositions of gasdermins, a listing of structural relatives of GSDMD^Cterm^ and an illustration of protein construct design and purification.

Supplemental_Figure1: Structures and purification of GSDMD^Cterm^, GSDMA3^FL^ and GSDMB^Cterm^

Supplemental_Figure2: Sequence alignment of all GSDM family members

Supplemental_Figure3: Sequence alignment of GSDMDs from different species

Supplemental_Figure4: Sequence alignment of GSDMD, GSDMA3 and GSDMB

supplemental_Table_I: PDBeFold search for structures related to GSDMD^Cterm^

## REFERENCES

1. Akira, S., S. Uematsu, and O. Takeuchi, Pathogen Recognition and Innate Immunity. Cell, 2006. 124: p. 783–801.

2. Schaefer, L., Complexity of Danger: The Diverse Nature of Damage-Associated Molecular Patterns. 2014. 289(51): p. 35237–35245.

3. Martinon, F., K. Burns, and J. Tschopp, The Inflammasome: A Molecular plattform Triggering Activation of Inflammatory caspases and processing of proIL-β. Molecular Cell, 2002. 10(2): p. 417–426.

4. Latz, E., T.S. Xiao, and A. Stutz, Activation and regulation of the inflammasomes. Nature Reviews Immunology, 2013. 13(6): p. 397–411.

5. Sollberger, G., et al., Caspase-1: The inflammaosme and beyond. Innate Immunity, 2014. 20(2): p. 115–125.

6. Miao, E., et al., Caspase-1 induced pyroptosis is an innate immune effector mechanism against intracellular bacteria. Nature Immunology, 2010. 11(12): p. 1136–1142.

7. Brodsky, I. and R. Medzhitov, Pyroptosis: macrophage suicide exposes hidden invaders. Current Biology, 2011. 21(2): p. R72–R75.

8. Shi, J., et al., Cleavage of GSDMD by inflammatory caspases determines pyroptotic cell death. Nature, 2015. 526(7575): p. 660–665.

9. Kayagaki, N., et al., Caspase-11 cleaves gasdermin D for non-canonical inflammasome signaling. Nature, 2015. 526(7575): p. 666–671.

10. Ding, J., et al., Pore-forming activity and structural autoinhibition of the gasdermin family. Nature, 2016. 535(7610): p. 111–116.

11. Aglietti, R.A., et al., GsdmD p30 elicited by caspase-11 during pyroptosis forms pores in membranes. PNAS, 2016. 113(28): p. 7858–7863.

12. Sborgi, L., et al., GSDMD membrane pore formation constitutes the mechanism of pyroptotic cell death. EMBO Journal, 2016. 35(16): p. 1766–78.

13. Liu, X., et al., Inflammasome-activated gasdermin D causes pyroptosis by forming membrane pores. Nature, 2016. 535(7610): p. 153–158.

14. Rogers, C., et al., Cleavage of DFNA5 by caspase-3 during apoptosis mediates progression to secondary necrotic/pyroptotic cell death. Nature Communications, 2017. 8: p. 14128.

15. Zihilf, M., et al., Association Between Gasdermin A and Gasdermin B Polymorphisms and Suseptibility to Adult and Childhood Asthma Among Jordanians. Genet Test Mol Biomarkers, 2016. 20(3): p. 143–8.

16. Du, M., et al., Prostate cancer risk locus at 8q24 as a regulatory hub by physical interactions with multiple genomic loci acros the genome. Hum Mol Genet., 2015. 24(1): p. 154–66.

17. Chao, K., L. Kulakova, and O. Herzberg, Gene Polymorphism Linked to Increased Asthma and IBD Risk Alters Gasdermin-B Structure, a Sulfatide and Phosphoinositide Binding protein. PNAS, 2017. 114(7): p. E1128–E1137.

18. Doerks, T., Systematic Identification of Novel Protein Domain Families Associated with Nuclear Functions. Genome Res., 2002. 12(1): p. 47–56.

19. Kabsch, W., XDS. Acta Crystallographica Section D, 2010. 66(2): p. 125–132.

20. Kabsch, W., Integration, scaling, space-group assignment and post-refinement. Acta Crystallographica Section D, 2010. 66(2): p. 133–144.

21. McCoy, A.J., et al., Phaser crystallographic software. Journal of Applied Crystallography, 2007. 40(4): p. 658–674.

22. Emsley, P., et al., Features and development of Coot. Acta Crystallographica Section D, 2010. 66(4): p. 486–501.

23. Zhang, K.Y.J., K. Cowtan, and P. Main, Combining constratins for electron density. Methods in Enzymology, 1997. 277: p. 53–64.

24. Cowtan, K., The Buccaneer software for automated model building. Acta Cryst., 2006. 62: p. 10002–1011.

25. Cowtan, K., Fitting molecular fragments into electron density. Acta Cryst., 2008. 64: p. 83–89.

26. Smart, O.S., et al., Exploiting structure similarity in refinement: automated NCS and target-structure restraints in BUSTER. Acta Cryst., 2012. 68(Pt 4): p. 368–380.

27. Afonine, P.V., et al., Automatic multiple-zone rigid-body refinement with a large convergence radius. Journal of Applied Crystallography, 2009. 42(4): p. 607–615.

28. Afonine, P.V., et al., Towards automated crystallographic structure refinement with phenix.refine. Acta Crystallographica Section D, 2012. 68(4): p. 352–367.

29. Afonine, P.V., et al., Bulk-solvent and overall scaling revisited: faster calculations, improved results. Acta Crystallographica Section D, 2013. 69(4): p. 625–634.

30. Headd, J.J., et al., Use of knowledge-based restraints in phenix.refine to improve macromolecular refinement at low resolution. Acta Crystallographica Section D, 2012. 68(4): p. 381–390.

31. Krissinel, E. and K. Henrick, Secondary-Structure matching (SSM), a new tool for fast protein structure alignment in three dimensions. Acta Cryst., 2004. 60(Pt 12 Pt 1): p. 2256–2268.

32. Krissinel, E. and K. Henrick, Inference of Macromolecular Assemblies from Crystalline State. Journal of Molecular Biology, 2007. 372(3): p. 774–797.

33. Sievers, F., et al., Fast, scalable generation of high-quality protein multiple sequence alignments using Clustal Omega. EMBO Report, 2011. 7: p. 539.

34. Goujon, M., et al., A new bioinformatics analysis toolsframework at EMBL-EBI. Nucleic Acids Research, 2010. 38: p. 695–699.

35. Pei, J. and N.V. Grishin, AL2CO: calculation of positional conservation in a protein sequence alignment. Bioinformatics., 2001. 17(8): p. 700–712.

